# Effects of Probiotics, Glucose Oxidase and its Combination on Immune Function, Antioxidant Status, Serum Biochemical Incides and Toxin Residue in Sheep when Challenged with Aflatoxin B1

**DOI:** 10.1101/2023.12.25.573291

**Authors:** Yu Zhang, Henan Lu, Erdene-Khas, Caixia Zhang, Hairong Wang

## Abstract

The objective of this study was to investigate the ability of a mixed mycotoxins detoxification agent (probiotics, Glucose oxidase (GOD) and its combination) to alleviate the toxicity of aflatoxin B1 (AFB1) by assessing growth performance, serum toxin residue, immune function, antioxidant status and serum biochemical indices of sheep. Twenty 8-month-old Mongolian sheep were randomly assigned to 1-5 diet treatment groups: 1) the control (CON) group: basal diet; 2) the AFB1 (A) group: CON + 0.2 mg/kg AFB1; 3) the AFB1 and probiotics (AP) group: A + 0.5% probiotics; 4) the AFB1 and GOD (AG) group: A + 0.5% GOD; 5) the AFB1, probiotics, and GOD (APG) group: A + 0.5% probiotics + 0.5% GOD. Results showed that treatment A adversely affected the health and performance of sheep. However, the treatments AP, AG, or APG exerted a positive effect on health, performance and other indications. In conclusion, probiotics, GOD, and its combination induced injury of Mongolian sheep by alleviating the negative effects of AFB1 on the production performance, immune indexes, antioxidant indexes, and serum biochemical indexes and effectively reducing serum toxin residue.

**Key Contribution:** The study investigated the effects of probiotics, Glucose oxidase and its combination on serum detoxification of AFB1 by assessing growth performance, serum toxin residue, immune function, antioxidant status and blood biochemical indices of sheep. AFB1 - contaminated diets supplemented with probiotics and GOD were effective in improving growth performance, immunity, antioxidant function, liver function, and decreasing serum AFB1 residue of sheep.

## 1. Introduction

Mycotoxins are secondary metabolites produced by pathogenic fungi species. In general, mycotoxins are typically easy to accumulate and lead to significant negative impacts on animals (Benjamin et al., 2020). Aflatoxins (AFs), as highly toxic mycotoxins, produced by fungi of the Aspergillus genus, especially A. flavus (Rushing and Selim, 2019). Among the Afs forms (AFB1, B2, G1, G2, and M1), aflatoxin B1 (AFB1) is the most common and the most potential form (Yhnly and Schwartz, 2020). International Agency for Research on Cancer (IARC) showed that AFB1 has an important association with the occurrence of hepatocellular carcinoma, which is clearly classified as the first class of carcinogens (Véronique et al., 2012). AFB1 causes animal organism health effects, which is commonly found in ruminant diets. For instance, it may results in reduced growth performance and feed conversion ratio (Edrington et al., 1994, Gallo et al., 2016), depress immune competence (Fernandez et al., 2000), create oxidative stress (Huang et al., 2018), inhibit liver function (B. et al., 1991) and influence reproductive performance (Komsky et al., 2018).

Physical, chemical, and biological methods adopted for the elimination of AFB1 are widely used. However, certain nutrients are destroyed in the physical process (Zolfaghari et al., 2020), chemical methods relate to food security and are restricted severely, because there may have chemical residues (Fernandez et al., 2000), along with unhealthy effects on humans. Biological detoxification measure is safer and more efficient than the other ways, especially probiotics and enzymes that have been utilized for their ability either absorb or degrade AFB1 (Gallo et al., 2016). In recent years, adding probiotics to animal diets has become a popular AFB1 detoxification method. Since probiotics can bind to AFs entry into the gastrointestinal tract, thus can block the absorption of mycotoxin by the organism and reduce the negative impacts on animal health (Gallo et al., 2016). Several species of probiotics have found that to reduce the residual level of AFB1 (Chen et al., 2022, Poloni et al., 2020, Sui-Sheng and James, 1999, Topcu et al., 2010, Wang et al., 2018), decrease oxidative damage (Xia et al., 2017, Li et al., 2021). A previous study by our research group has demonstrated that these probiotics degrade AFB1 with well degradation efficiency in vitro (Peng, 2018). However, despite the previous finding of mono-species probiotics can degrade AFB1, it remains unknown whether multi-species degrade AFB1 has a synergetic effect and relevant studies are rare.

Glucose oxidase (GOD) is an aerobic dehydrogenase that catalyzes the oxidation of β-d-glucose into gluconic acid and hydrogen peroxide (Bankar et al., 2009). Such enzyme is already widely used in the feed production industry since it has been established that GOD has effects on bacteriostatic effect (Sandholm et al., 1988), growth-promotion (Wu et al., 2019), improving antioxidant capacity (Dang et al., 2022, Zhang et al., 2020), maintaining intestinal microbiota homeostasis (Qu and Liu, 2021). Furthermore, hydrogen peroxide produced during the GOD redox process can detoxify mycotoxins, especially AFB1 which contains a xanthene phosphorous copper group through oxidative dehydrogenation (Murphy K et al., 2008). However, GOD as a promising AFB1 toxic inhibitor has not been well investigated on the effects in counteracting the growth inhibition, hepatic toxicity, immune toxicity and toxic residues in ruminants induced by AFB1, besides, there are rarely relevant studies about it.

Thus, combine various lines of evidence that supported the ingredients in this experiment, the objective of this research is to explore the influences of Mongolian sheep fed with AFB1-contaminated diets supplemented with probiotics and GOD on growth performance, immune response, antioxidative status, serum biochemistry profiles, and Serum toxin residues, to evaluate the effects of adding probiotics and GOD in vivo on sheep fed with AFB1-contaminated diet. The result of the study will provide a theoretical basis for further development of a detoxication agent for AFB1.

## 2. Results

### 2.1 Growth performance

The effects of probiotics and GOD on growth performance in sheep feeding with AFB1 are reported in Table 2. During 1 to 14 d, the ADG in group A, and APG were significantly lower than other three groups (P < 0.05), the ADFI in group C was significantly higher than other test groups (P < 0.05). There were no difference among all treatments in ADFI.

During 15 to 28 d, compared to group C, ADG in group A significantly decreased by 40% (P < 0.05), but on significant difference in other test groups (P > 0.05). In addition, there was a trend toward higher ADFI in the group AG compared with A (P < 0.10).

Throughout the entire trail (1 to 28 d), group A had a significant decreased in ADG, and ADFI (P < 0.05) but significantly increased F/G compared to C group (P < 0.05). AP, AG, and APG significantly increased ADG (P < 0.05), decreased F/G than group A (P < 0.05), no differences were observed in ADFI (P > 0.05).

**Table. 1.**
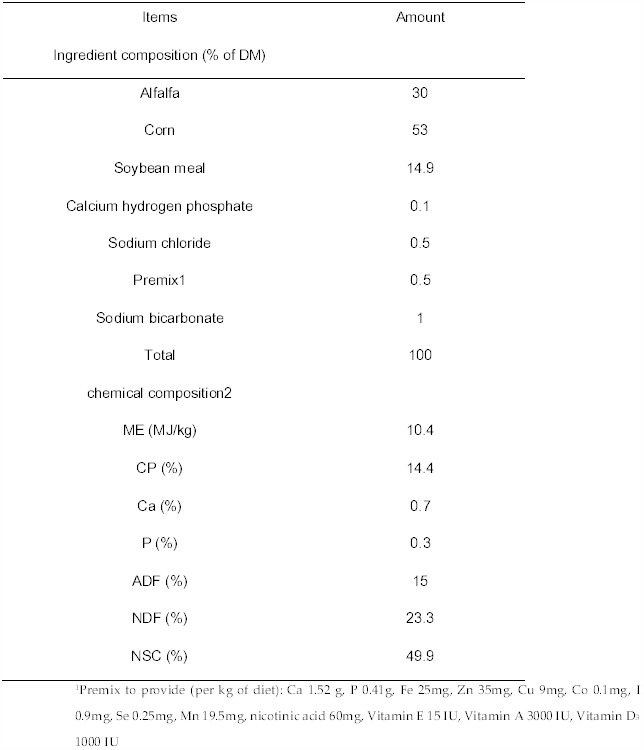
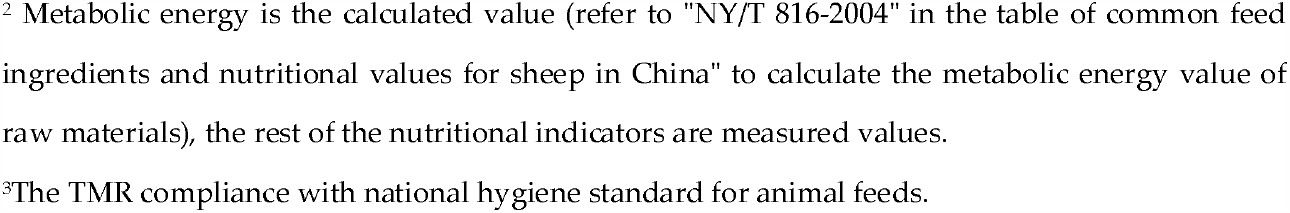
Ingredients and chemical composition of the basal diet.

| Items | Amount |
| --- | --- |
| Ingredient composition (% of DM) |  |
| Alfalfa | 30 |
| Corn | 53 |
| Soybean meal | 14.9 |
| Calcium hydrogen phosphate | 0.1 |
| Sodium chloride | 0.5 |
| Premix<sup>1</sup> | 0.5 |
| Sodium bicarbonate | 1 |
| Total | 100 |
| chemical composition<sup>2</sup> |  |
| ME (MJ/kg) | 10.4 |
| CP (%) | 14.4 |
| Ca (%) | 0.7 |
| P (%) | 0.3 |
| ADF (%) | 15 |
| NDF (%) | 23.3 |
| NSC (%) | 49.9 |

<sup>1</sup>Premix to provide (per kg of diet): Ca 1.52 g, P 0.41g, Fe 25mg, Zn 35mg, Cu 9mg, Co 0.1mg, I 0.9mg, Se 0.25mg, Mn 19.5mg, nicotinic acid 60mg, Vitamin E 15 IU, Vitamin A 3000 IU, Vitamin D<sub>3</sub> 1000 IU
<sup>2</sup> Metabolic energy is the calculated value (refer to "NY/T 816-2004" in the table of common feed ingredients and nutritional values for sheep in China" to calculate the metabolic energy value of raw materials), the rest of the nutritional indicators are measured values.
<sup>3</sup>The TMR compliance with national hygiene standard for animal feeds.

**Table 2.**
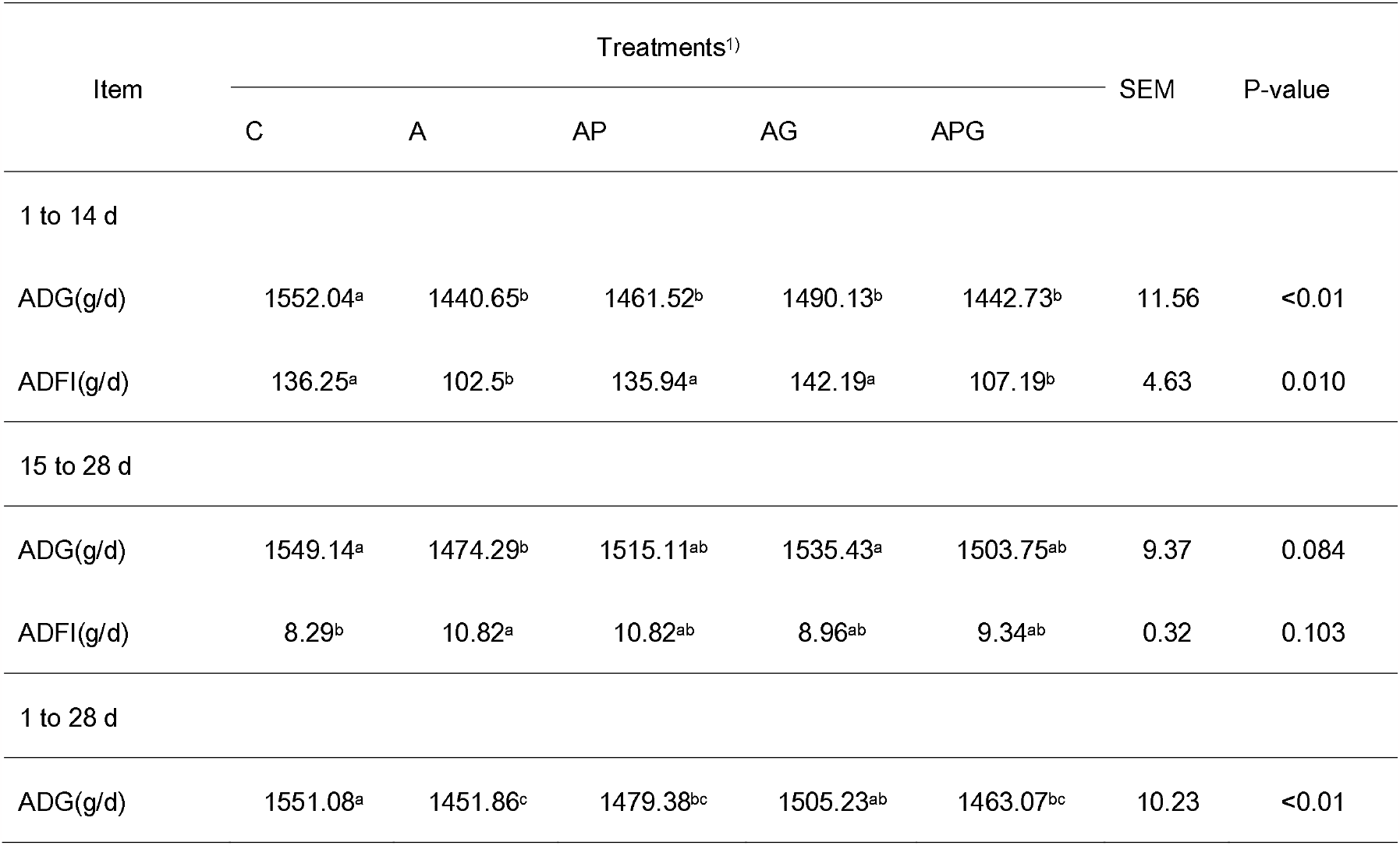
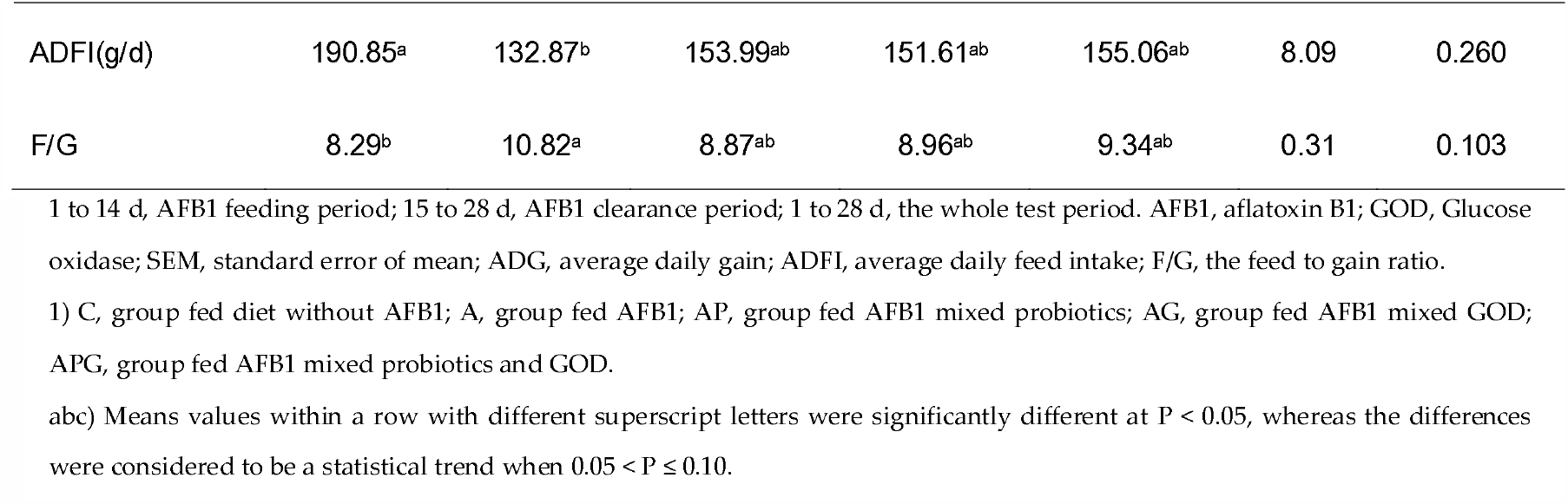
Effect of probiotics and Glucose oxidase on growth performance in sheep challenged with AFB1.

| Item | Treatments<sup>1)</sup> |  |  |  |  | SEM | P-value |
| --- | --- | --- | --- | --- | --- | --- | --- |
|  | C | A | AP | AG | APG |  |  |
| 1 to 14 d |  |  |  |  |  |  |  |
| ADG(g/d) | 1552.04<sup>a</sup> | 1440.65<sup>b</sup> | 1461.52<sup>b</sup> | 1490.13<sup>b</sup> | 1442.73<sup>b</sup> | 11.56 | <0.01 |
| ADFI(g/d) | 136.25<sup>a</sup> | 102.5<sup>b</sup> | 135.94<sup>a</sup> | 142.19<sup>a</sup> | 107.19<sup>b</sup> | 4.63 | 0.010 |
| 15 to 28 d |  |  |  |  |  |  |  |
| ADG(g/d) | 1549.14<sup>a</sup> | 1474.29<sup>b</sup> | 1515.11<sup>ab</sup> | 1535.43<sup>a</sup> | 1503.75<sup>ab</sup> | 9.37 | 0.084 |
| ADFI(g/d) | 8.29<sup>b</sup> | 10.82<sup>a</sup> | 10.82<sup>ab</sup> | 8.96<sup>ab</sup> | 9.34<sup>ab</sup> | 0.32 | 0.103 |
| 1 to 28 d |  |  |  |  |  |  |  |
| ADG(g/d) | 1551.08<sup>a</sup> | 1451.86<sup>c</sup> | 1479.38<sup>bc</sup> | 1505.23<sup>ab</sup> | 1463.07<sup>bc</sup> | 10.23 | <0.01 |
| ADFI(g/d) | 190.85<sup>a</sup> | 132.87<sup>b</sup> | 153.99<sup>ab</sup> | 151.61<sup>ab</sup> | 155.06<sup>ab</sup> | 8.09 | 0.260 |
| F/G | 8.29<sup>b</sup> | 10.82<sup>a</sup> | 8.87<sup>ab</sup> | 8.96<sup>ab</sup> | 9.34<sup>ab</sup> | 0.31 | 0.103 |

1 to 14 d, AFB1 feeding period; 15 to 28 d, AFB1 clearance period; 1 to 28 d, the whole test period. AFB1, aflatoxin B1; GOD, Glucose oxidase; SEM, standard error of mean; ADG, average daily gain; ADFI, average daily feed intake; F/G, the feed to gain ratio. 1) C, group fed diet without AFB1; A, group fed AFB1; AP, group fed AFB1 mixed probiotics; AG, group fed AFB1 mixed GOD; APG, group fed AFB1 mixed probiotics and GOD. abc) Means values within a row with different superscript letters were significantly different at $P < 0.05$ , whereas the differences were considered to be a statistical trend when $0.05 < P \leq 0.10$ .

### 3.2 Serum toxin residue

On d 14, fed the AFB-contaminated diets increased (P < 0.05) serum toxin residue level, but AG or APG supplementation of AFB-contaminated diets decreased it (P < 0.01) (Table 3).On d 21, the addition of an AFB-contaminated diet significantly increased (P < 0.05) serum toxin residue, but adding AP and APG dramatically decreased it (P < 0.05).On d 28, addition of AP, AG, and APG did not significantly affect the serum toxin residue.

**Table 3.**
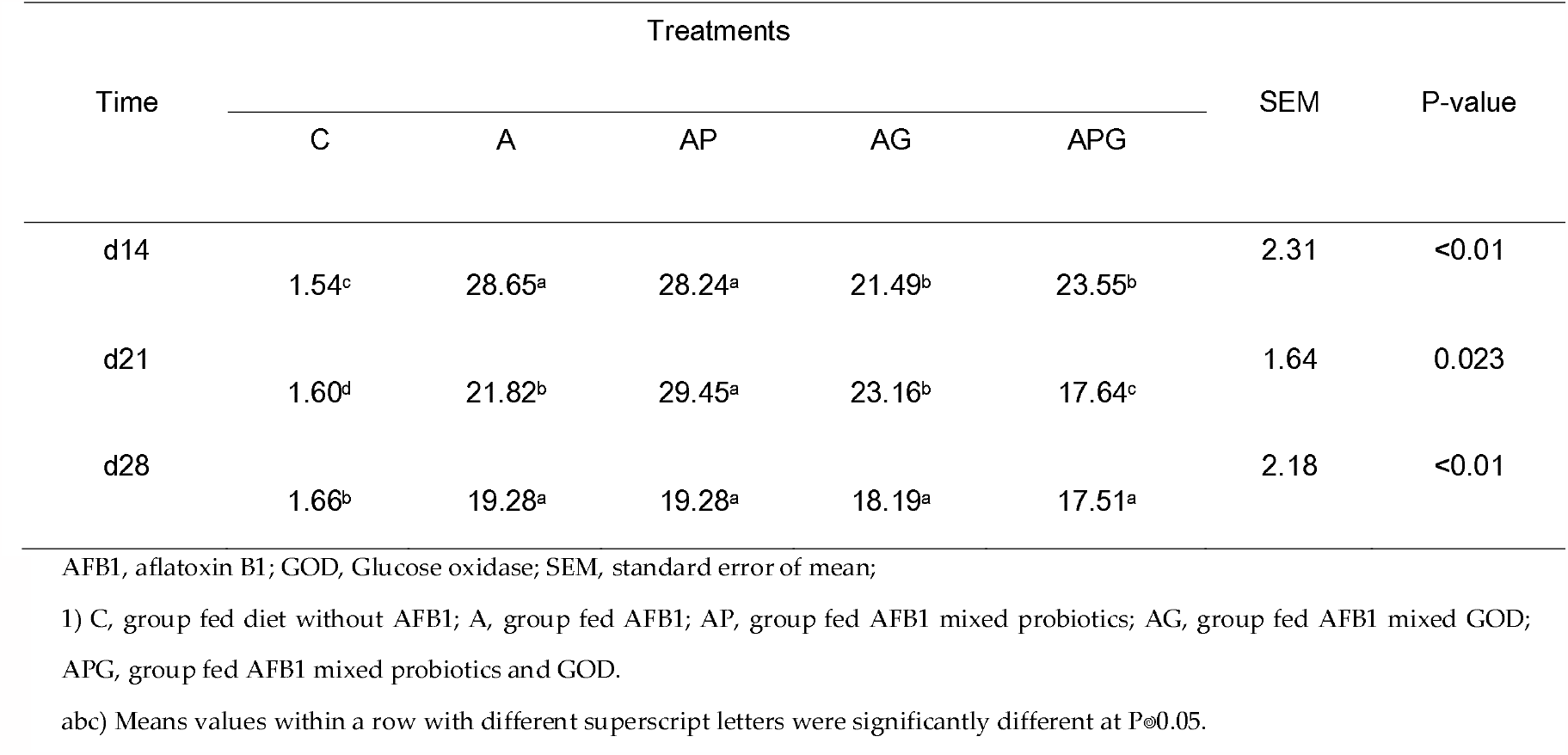
Effects of probiotics and Glucose oxidase on serum toxin residue in sheep feeding with AFB1 (μg/L)

| Time | Treatments |  |  |  |  | SEM | P-value |
| --- | --- | --- | --- | --- | --- | --- | --- |
|  | C | A | AP | AG | APG |  |  |
| d14 | 1.54<sup>c</sup> | 28.65<sup>a</sup> | 28.24<sup>a</sup> | 21.49<sup>b</sup> | 23.55<sup>b</sup> | 2.31 | <0.01 |
| d21 | 1.60<sup>d</sup> | 21.82<sup>b</sup> | 29.45<sup>a</sup> | 23.16<sup>b</sup> | 17.64<sup>c</sup> | 1.64 | 0.023 |
| d28 | 1.66<sup>b</sup> | 19.28<sup>a</sup> | 19.28<sup>a</sup> | 18.19<sup>a</sup> | 17.51<sup>a</sup> | 2.18 | <0.01 |

AFB1, aflatoxin B1; GOD, Glucose oxidase; SEM, standard error of mean; 1) C, group fed diet without AFB1; A, group fed AFB1; AP, group fed AFB1 mixed probiotics; AG, group fed AFB1 mixed GOD; APG, group fed AFB1 mixed probiotics and GOD. abc) Means values within a row with different superscript letters were significantly different at $P \leq 0.05$ .

### 3.3 Immunity Parameter

Immunity indices of sheep are given at Table 4. On d 14, no significant effects of dietary AFB1 supplementation on immunity indices (P > 0.05). Compared to the A, AP significantly increased serum IgM (P < 0.05), AG markedly increased IL-2, and IgG (P < 0.05). Compared to the AP, and AG, IL-2 in APG significantly decreased (P < 0.05),IL-6, and IL-10 level tend to be remarkably decreased (0.05 < P < 0.1). The serum TNF-α level in APG was less than AG (0.05 < P < 0.1).

**Table 4.**
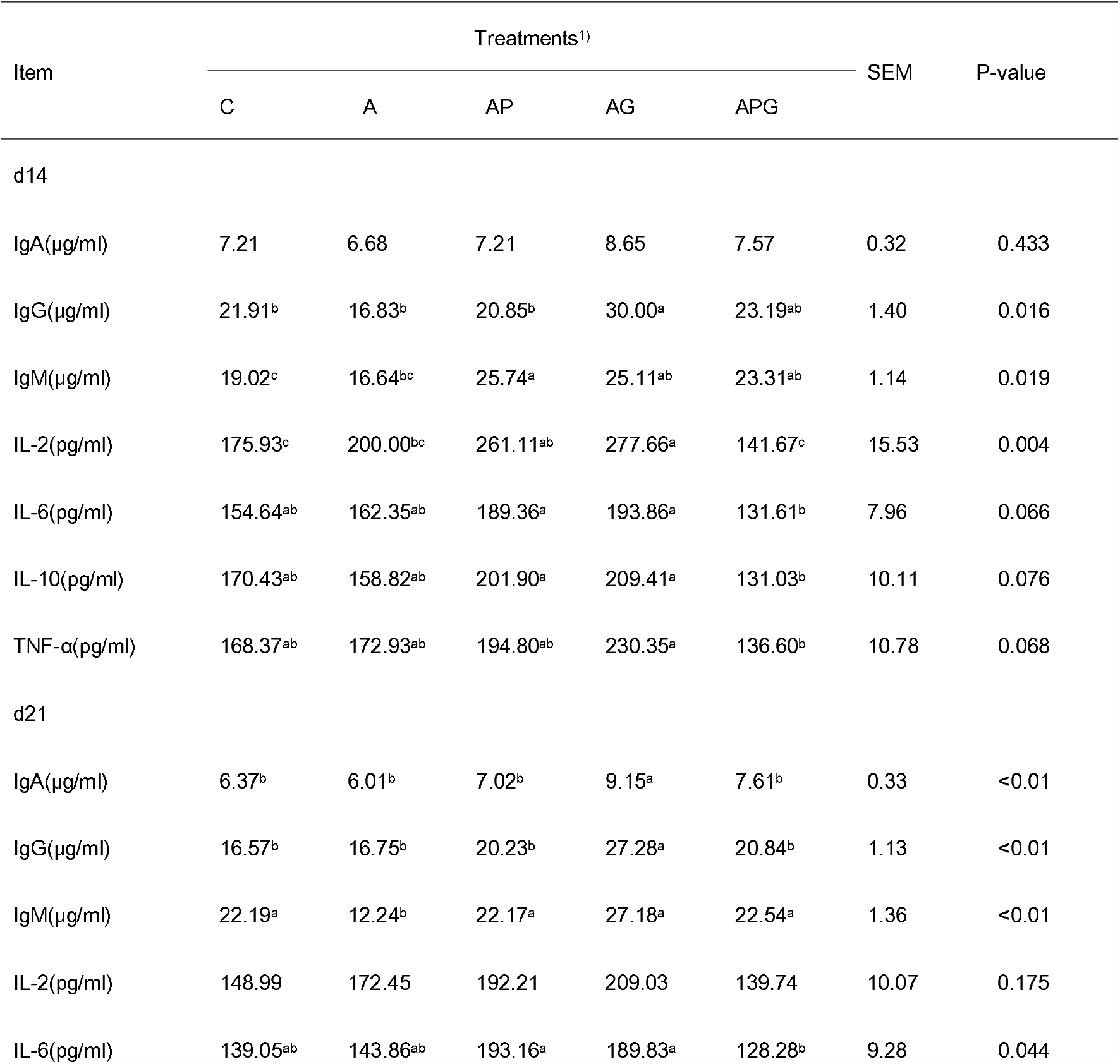
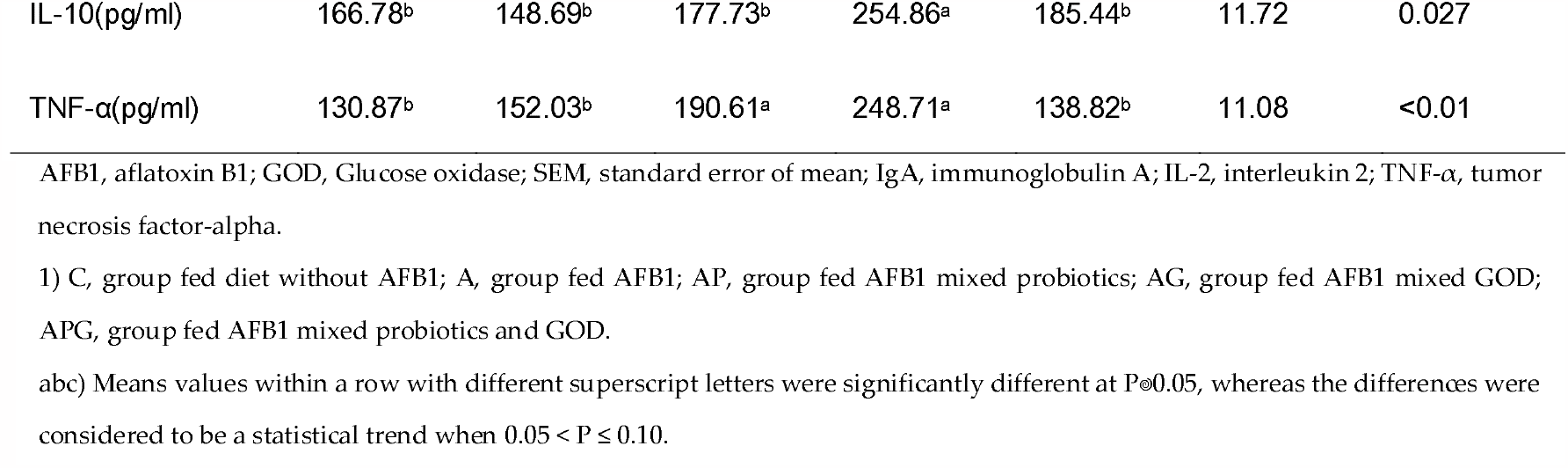
Effects of probiotics and Glucose oxidase on Immunity parameters in sheep challenged with AFB1.

On d 21, compared to the C, A significantly decreased IgM (P < 0.05). compared to the A, IgA, and IgG level in AG significantly increased (P < 0.05), IgM level in AP, AG, and APG significantly increased (P < 0.05). Compared to the AP, and AG, APG significantly decreased serum IL-6, and TNF-α level (P < 0.05), but, APG had no significant difference in IL-6, and TNF-α level than control group (P > 0.05). IL-10 level in the AG group was higher than other groups (P < 0.05).

### 3.4 Antioxidant capacity

Table 5 summarized the antioxidant status of sheep. On d 14, sheep fed AFB-contaminated diets recorded a significantly lower T-AOC, CAT, and SOD activity (P < 0.05). Compared to the A, the addition of AP or APG resulted in a significantly increased T-AOC and SOD activity (P < 0.05). AP, AG, and APG supplementation increased CAT activity (P < 0.05). Moreover, the addition of AP or AG had increasing GSH-Px activity by 71.46% and 82.22% (P < 0.05). On d 21, sheep fed AFB-contaminated diets had significantly lower T-AOC activity than the control group (P < 0.05). Compared to the A, AG significantly increased serum GSH-Px activity (P < 0.05). SOD activity in AP, AG, or APG tend to be remarkably increased (0.05 < P < 0.1), AP or APG significantly reduced serum MDA level (P < 0.05).

**Table 5.** Effects of probiotics and Glucose oxidase on antioxidant status in sheep feeding with AFB1.

| Item | Treatments<sup>1)</sup> |  |  |  |  | SEM | P-value |
| --- | --- | --- | --- | --- | --- | --- | --- |
|  | C | A | AP | AG | APG |  |  |
| d14 |  |  |  |  |  |  |  |
| T-AOC(U/ml) | 0.56<sup>a</sup> | 0.46<sup>b</sup> | 0.54<sup>a</sup> | 0.51<sup>ab</sup> | 0.55<sup>a</sup> | 0.01 | 0.048 |
| CAT(U/ml) | 60.75<sup>a</sup> | 41.95<sup>b</sup> | 61.33<sup>a</sup> | 63.89<sup>a</sup> | 72.79<sup>a</sup> | 3.19 | 0.011 |
| GSH-Px(ng/ml) | 11.69<sup>ab</sup> | 8.27<sup>bc</sup> | 14.18<sup>a</sup> | 15.07<sup>a</sup> | 5.69<sup>c</sup> | 1.09 | 0.012 |
| SOD(U/ml) | 22.05<sup>a</sup> | 17.62<sup>b</sup> | 22.52<sup>a</sup> | 20.42<sup>ab</sup> | 22.23<sup>a</sup> | 0.55 | 0.021 |
| MDA(nmol/ml) | 1.91 | 2.33 | 2.18 | 2.08 | 1.93 | 0.09 | 0.693 |
| d21 |  |  |  |  |  |  |  |
| T-AOC(U/ml) | 0.57<sup>a</sup> | 0.48<sup>b</sup> | 0.46<sup>b</sup> | 0.51<sup>b</sup> | 0.51<sup>b</sup> | 0.01 | <0.01 |
| CAT(U/ml) | 63.54 | 53.88 | 59.62 | 62.36 | 67.53 | 1.71 | 0.116 |
| GSH-Px(ng/ml) | 10.87<sup>b</sup> | 7.61<sup>b</sup> | 10.33<sup>b</sup> | 16.44<sup>a</sup> | 8.28<sup>b</sup> | 0.79 | <0.01 |
| SOD(U/ml) | 22.17<sup>ab</sup> | 19.90<sup>b</sup> | 25.09<sup>a</sup> | 23.84<sup>a</sup> | 23.72<sup>a</sup> | 0.63 | 0.066 |
| MDA(nmol/ml) | 1.85<sup>ab</sup> | 2.28<sup>a</sup> | 1.68<sup>b</sup> | 2.23<sup>a</sup> | 1.58<sup>b</sup> | 0.09 | 0.014 |

1) C, group fed diet without AFB1; A, group fed AFB1; AP, group fed AFB1 mixed probiotics; AG, group fed AFB1 mixed GOD; APG, group fed AFB1 mixed probiotics and GOD.
Means values within a row with different superscript letters were significantly different at $P \leq 0.05$ , whereas the differences were considered to be a statistical trend when $0.05 < p \leq 0.10$ .

### 3.5 Serum biochemical parameters

Data for the serum biochemical parameters of sheep are presented in Table 6. On d 14, sheep fed AFB-contaminated diets had significantly increased TBIL and DBIL levels (P < 0.05), whereas significantly decreased ALB (P < 0.05), tends to decrease TP contents (0.05 < P < 0.1). Compared to the A, ALT, and DBIL levels in AP, AG and APG significantly decreased (P < 0.05), the addition of AP resulted in a significantly decreased TBIL (P < 0.05).

**Table 6.**
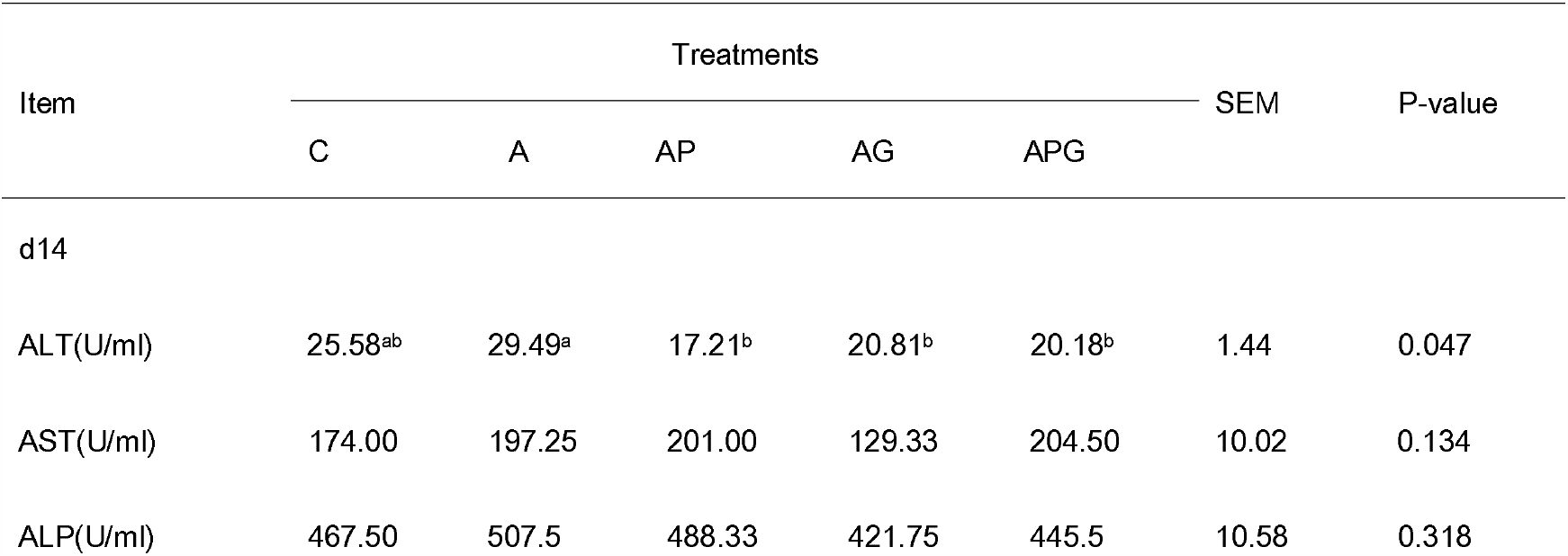
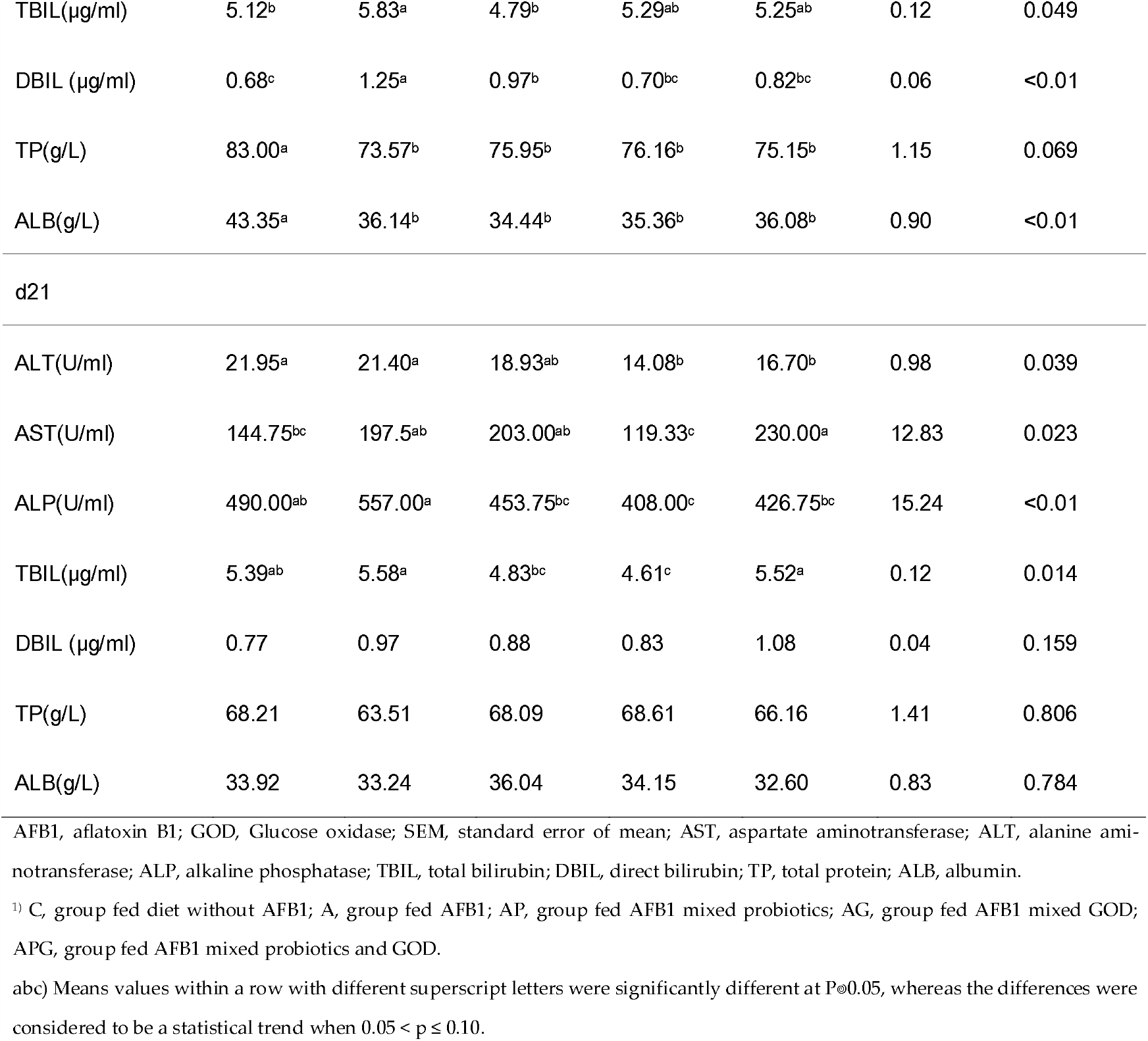
Effects of probiotics and Glucose oxidase on blood biochemical parameters in sheep feeding with AFB1.

| Item | Treatments |  |  |  |  | SEM | P-value |
| --- | --- | --- | --- | --- | --- | --- | --- |
|  | C | A | AP | AG | APG |  |  |
| d14 |  |  |  |  |  |  |  |
| ALT(U/ml) | 25.58<sup>ab</sup> | 29.49<sup>a</sup> | 17.21<sup>b</sup> | 20.81<sup>b</sup> | 20.18<sup>b</sup> | 1.44 | 0.047 |
| AST(U/ml) | 174.00 | 197.25 | 201.00 | 129.33 | 204.50 | 10.02 | 0.134 |
| ALP(U/ml) | 467.50 | 507.5 | 488.33 | 421.75 | 445.5 | 10.58 | 0.318 |
| TBIL(μg/ml) | 5.12<sup>b</sup> | 5.83<sup>a</sup> | 4.79<sup>b</sup> | 5.29<sup>ab</sup> | 5.25<sup>ab</sup> | 0.12 | 0.049 |
| DBIL (μg/ml) | 0.68<sup>c</sup> | 1.25<sup>a</sup> | 0.97<sup>b</sup> | 0.70<sup>bc</sup> | 0.82<sup>bc</sup> | 0.06 | <0.01 |
| TP(g/L) | 83.00<sup>a</sup> | 73.57<sup>b</sup> | 75.95<sup>b</sup> | 76.16<sup>b</sup> | 75.15<sup>b</sup> | 1.15 | 0.069 |
| ALB(g/L) | 43.35<sup>a</sup> | 36.14<sup>b</sup> | 34.44<sup>b</sup> | 35.36<sup>b</sup> | 36.08<sup>b</sup> | 0.90 | <0.01 |
| d21 |  |  |  |  |  |  |  |
| ALT(U/ml) | 21.95<sup>a</sup> | 21.40<sup>a</sup> | 18.93<sup>ab</sup> | 14.08<sup>b</sup> | 16.70<sup>b</sup> | 0.98 | 0.039 |
| AST(U/ml) | 144.75<sup>bc</sup> | 197.5<sup>ab</sup> | 203.00<sup>ab</sup> | 119.33<sup>c</sup> | 230.00<sup>a</sup> | 12.83 | 0.023 |
| ALP(U/ml) | 490.00<sup>ab</sup> | 557.00<sup>a</sup> | 453.75<sup>bc</sup> | 408.00<sup>c</sup> | 426.75<sup>bc</sup> | 15.24 | <0.01 |
| TBIL(μg/ml) | 5.39<sup>ab</sup> | 5.58<sup>a</sup> | 4.83<sup>bc</sup> | 4.61<sup>c</sup> | 5.52<sup>a</sup> | 0.12 | 0.014 |
| DBIL (μg/ml) | 0.77 | 0.97 | 0.88 | 0.83 | 1.08 | 0.04 | 0.159 |
| TP(g/L) | 68.21 | 63.51 | 68.09 | 68.61 | 66.16 | 1.41 | 0.806 |
| ALB(g/L) | 33.92 | 33.24 | 36.04 | 34.15 | 32.60 | 0.83 | 0.784 |

AFB1, aflatoxin B1; GOD, Glucose oxidase; SEM, standard error of mean; AST, aspartate aminotransferase; ALT, alanine aminotransferase; ALP, alkaline phosphatase; TBIL, total bilirubin; DBIL, direct bilirubin; TP, total protein; ALB, albumin.
<sup>1)</sup> C, group fed diet without AFB1; A, group fed AFB1; AP, group fed AFB1 mixed probiotics; AG, group fed AFB1 mixed GOD; APG, group fed AFB1 mixed probiotics and GOD.
abc) Means values within a row with different superscript letters were significantly different at $P \leq 0.05$ , whereas the differences were considered to be a statistical trend when $0.05 < p \leq 0.10$ .

On d 21, there were no significant changes in serum biochemical parameters of sheep feed AFB-contaminated diets. Compared to the A, the addition of AG or APG significantly decreased ALT activity (P < 0.05), AG significantly decreased serum AST activity (P < 0.05), AP, AG or APG had significant lower ALP activity (P < 0.05). Reduction in TBIL level in sheep serum by added AP or AG (P < 0.05).

## 3 Discussion

AFB1, a highly toxic carcinogen, is a powerful mutagen that could decrease feed conversion efficiency, cause diseases, improve mortality rate and result in enormous economic losses worldwide. In a previous study, average daily gain, feed intake and feed to gain ratio (F / G) were decreased among lambs that were fed 2.5 mg of AFB1 / kg diet (Deng et al., 2020). Such adverse effects may have resulted from anorexia, listlessness and the inhibitory effects of AFB1 on protein synthesis and lipogenesis, probably as a consequence of a deterioration of the digestive and metabolic efficiency (Fernandez et al., 2000, Oguz and Kurtoglu, 2000). Similar as the previous one, the present study showed that average daily gain, and feed intake were depressed in sheep fed AFB1, the feed to gain ratio (F / G) of the whole experiment was elevated. Taken together, this suggests that the effects of AFB1 on sheep growth performance might be related to the decrease of feed intake. The addition of absorbents to the AFB-contaminated diets did not affect dry matter intake (Kutz et al., 2009, Queiroz et al., 2012). The results of this study are in line with the previous ones, probiotics, and glucose oxidase had no significant effects on the average daily feed intake of sheep that fed AFB1 in the whole experiment. Bruno et al. showed that bacillus supplementation counteracts AFB1 negative effects on growth performance (Solis-Cruz et al., 2019). Dang et al. found that the enhancement of average daily gain of piglets could be improved by adding GOD (Dang et al., 2022). In our present study, probiotics and GOD could improve the average daily gain of sheep to different degrees during the entire trial, and feed to gain ratio (F / G) decreased significantly, some recovery ability in growth performance was shown through the way that adding two additives in AFB1 - contaminated diets, also detoxification was illustrated.

Serum cytokines and immunoglobulins levels could show the immunotoxicity of toxin tests, which were identified as valuable biological markers. With the inhibition of cellular immunity and humoral immunity, aflatoxin could decrease disease resistance, thus people and animals are more susceptible to infectious diseases (Huang et al., 2018, REDDY et al., 1987). Immunoglobulins are important immune factors, through which the B-cell lymphatic system recognize and defend the organism against specific pathogens or foreign materials (Murphy K et al., 2008). Previous research suggested that AFB1 may affect the immunocompetent of animal organisms (Ul-Hassan et al., 2012, Wang et al., 2019). Short - term exposure to AFB1 may exert immunosuppressive effects by inhibiting inflammation gene expression, while long-term exposure may upregulate the expression of proinflammatory cytokines to increase inflammatory response and apoptosis (Qian et al., 2014). Similar results were noted by Long et.al. (2016), the contents of serum IL - 6 and other inflammatory factors of mice were increased by performing the AFB1 challenge (Long et al., 2016). These are similar to the present results, where it was shown that the intake of AFB1 decreased immune-globulin concentration significantly and increased the content of IL - 6. The decrease of immunoglobulin content may be due to the inhibition of RNA polymerase activity after AFB1 enters the animal body, thus affecting the process of protein synthesis and reducing the amount of specific immunoglobulin synthesis (PIER, 1986). However, after adding probiotics and GOD, the immunoglobulin content in sheep serum was significantly increased. No apparent effects on the reduction of proinflammatory factor concentration when using probiotics and GOD. The rise of IL - 2, IL – 6, and TNF-α contents during this study, may relate to the stronger immune defense responses of the animal bodies that are stimulated by the elevation of proinflammatory cytokines doses (Wang et al., 2018). IL - 10, an immune regulatory cytokine, have inhibit effects on the synthesis of inflammatory cytokines, it works by adjusting the proliferation and differentiation of immune cells, also, it could prevent pathogens from entering bodies to help protect bodies. The increase of IL - 10 contents (d 21) showed that the body can strengthen the immunoregulatory ability by suppressing local and systematic inflammation, and in turn relieves inflammation.

Animal organisms under prolonged stress led to chronically continual that created reactive oxygen species (ROS) in their bodies that can cause cellular damage and oxidative damage to the body, which lead to oxidative stress (Fang et al., 2009). Oxidative stress is induced by mycotoxins, which can spur the production of ROS and destroy the antioxidant capacity of bodies even in a small dose (Reverberi et al., 2006). The antioxidant indicators T - AOC, CAT, GSH - Px, SOD, and MDA are the most important. The activity of T - AOC serves as an integrated parameter that reflects the overall antioxidant level of antioxidants and antioxidant enzymes. Results of previous studies have shown that AFB1 added to the animal’s diet will decrease the activities of T - AOC, GSH - Px, and SOD in serum, while increasing the MDA content (Huang et al., 2018, Rajput et al., 2017). The results of this study are in line with the previous ones, the T - AOC, CAT, GSH - Px, and SOD activities in the sheep serum were decreased to varying degrees. In contrast, sheep fed AFB1 - contaminated diets supplemented with probiotics and GOD showed a less oxidative stress level than AFB1 treated sheep. It might be observed these two additives increased the serum antioxidant enzymes activity, the reason may be antioxidant enzymes play an important role in decreases ROS production or inhibit the lipid peroxidation process resulted by AFB1 exposure (Naaz et al., 2014, Shen et al., 1996).

Blood biochemical parameters can reflect the nutritional metabolism condition of the animal body and health status. When the liver was threatened by Aflatoxin, liver cells were injured and increased membrane permeability, liver-specific enzyme (e.g., ALT, AST, and ALP) released into the bloodstream and leading to increased serum enzyme activity (Rajput et al., 2017, Yang et al., 2012). When hepatocytes are damaged, the ability to uptake, metabolism, and excrete bilirubin will decrease, this probably causes elevated serum bilirubin levels (Denli et al., 2009). In this study, compared with the control group, ALT, AST, and ALP activity of AFB1 - contaminated diets supplemented was increased, though the difference was not significant. In addition, feeding AFB1 - contaminated diets decreased ALB content and increased TBIL, and DBIL content on d 21. These parameters are the marker of liver damage, the increase of serum ALT, AST, ALP, TBIL and DBIL activity indicate that sheep liver might be impaired by AFB1 (Liu et al., 2020). The decrease in the content of ALB in serum might be because toxin inhibits the protein synthesis process, thereby damaging liver function. It was recently shown that exposure to AFB1 can rising ALT, AST, ALP, TBIL, and DBIL in rats, suggesting the hepatotoxicity of aflatoxin (Deng et al., 2020), which was consistent with the present study. In contrast, adding probiotics, GOD, or its combination in AFB1 - contaminated diets decreased those five indexes, suggesting that probiotics and GOD could abolish the adverse AFB1 effects on the liver function of sheep.

It is possible for AFB1 to be transmitted from livestock products to humans through food chains, posing a potential threat to human health. In this study, fed AFB1 - contaminated diets increase the toxin residue in sheep. As a result of oral administration, AFB1 is first absorbed into gastrointestinal track and then distributed to various tissues and body fluids throughout the body (Denli et al., 2009, Richard et al., 1983), but individual differences in aflatoxin residue and complete degradation time existed among the animals (Magnoli et al., 2011). It reported that probiotic supplement can markedly decrease aflatoxin residues in the liver as well as alleviate the negative effects of AFB1 (Salem et al., 2018). At the same time, the enzyme is also a good method of aflatoxin degradation. In this study, the AFB-contaminated diets supplemented with both probiotics and GOD exerted a significant beneficial effect in reducing AFB1 residue. It is speculated that low AFB1 residues by probiotics addition is owed to aflatoxin absorption and bio-degradation effect by the beneficial bacteria (Hernandez et al., 2009). In the case of this study, GOD reduced AFB1 residue level, probably because GOD disrupts the basis structure of AFB1 and resulted in a complete loss of its activity (Denli et al., 2009, Wang et al., 2019).

## 4 Conclusions

In conclusion, AFB1 - contaminated diets supplemented with probiotics and GOD were effective in improving growth performance, immunity, antioxidant function, liver function, and decreasing serum AFB1 residue of sheep. The combination of probiotics and GOD showed a better effect on improving immunity, and antioxidant function, reducing serum AFB1 residue than probiotics or GOD alone.

## 5 Materials and Methods

All laboratory animal studies were conducted in accordance with Animal Care and Use guidelines and were approved by the Institutional Animal Care and Use Committee at Inner Mongolia Agricultural University.

### 2.1 Experimental materials

The standard AFB1 (purity >99%) used for this experiment was purchased from Yuduo Bio-Technology Co., Ltd (Shanghai, China), and GOD was purchased from Yuanye Bio-Technology Co., Ltd. (Shanghai, China). The probiotics used contained Saccharomyces cerevisiae, Candida krusei, Bacillus amyloliquefaciens, Baclicus lincheniformis, Enterococcus faecalis, Lactobacillus fementum, Bacillus subtilis, those probiotics were mixed. Previous research of our research group found that Saccharomyces cerevisiae exhibited best degradation efficacy, the dose was doubled. The number of viable bacteria was determined by a plate count (final result 1.4 × 1010 CFU/g). These strains were kindly provided by the Animal Nutrition Laboratory of Inner Mongolia Agricultural University.

### 2.2 Experimental Design and Diets

The experiment was conducted at Inner Mongolia Agricultural University. Twenty 8-months-old Mongolian sheep were selected and randomly divided into five treatment groups (n = 4), which fed for 35 days. Treatments included the following: (1) basal diet (C), (2) AFB1 diet (A) containing C received 0.2 mg of AFB1/kg of the diet dry matter (DM), (3) AFB1-probiotics diet (AP) containing A with 0.5% probiotics of the diet DM, (4) AFB1-GOD diet (AG) containing A received 0.5% GOD of the diet DM, (5) AFB1-probiotics-GOD diet (APG) containing A with 0.5% probiotics, and 0.5% GOD of the diet DM. Body weight and body weight gain were calculated daily, the blood samples we collected from the jugular veins on day14, 21, and 28. The standard AFB1 was dissolved in methanol. An acclimation period of 7 days were allowed for sheep adaptation to the pen, followed by AFB1 was added for 14 days (d1 to 14), and finally clearance for 14 days (d15 to 28).

Diets were formulated according to nutrient requirements (Qu and Liu, 2021) for sheep. The ingredient composition and nutrient content of the basal diet is given in Table 1. The daily doses of AFB1, probiotics, and Glucose oxidase were divided into two aliquots, and each aliquot was mixed with 100 g of corn meal to encourage complete consumption. The AFB1 and two additive mixtures were fed to sheep within a separate container. The TMR was not provided until the mixture completely consumed to ensure sheep eat all of the AFB1 and additives (Queiroz et al., 2012). The control sheep were fed 100 g of corn meal before providing TMR at each feeding. Sheep were housed in a single column, to ensure a comfortable environment, and orderly cleaning for individual feeding to minimize stress responses, and fed with TMR at 0600 and 1700 h. Sheep were weighed at the beginning and end of each period before the morning feeding. Feed intake was recorded daily, and the gain to feed ratio was calculated.

### 2.3 Sample and data collection

Blood samples (10 mL) were collected from the jugular vein before morning feed on d 14, 21, and 28 respectively, and then centrifuged at 3000 × g for 15 min to obtain serum which was stored at last −20°C until later analysis. Immune parameters, including immunoglobulin A (IgA), immunoglobulin G (IgG), immunoglobulin M (IgM), interleukin 2 (IL-2), interleukin 6 (IL-6), interleukin 10 (IL-10), tumor necrosis factor-alpha (TNF-α) were analyzed with ELISA kits (Jiyinmei Biological Technology Co., LTD, Wuhan, China). Antioxidant status, including total anti-oxidant capacity (T-AOC), catalase (CAT), glutathione peroxide (GSH-Px), superoxide dismutase (SOD), malondialdehyde (MDA) were analyzed with assay kits (Solarbio Science & Technology Co Ltd. Beijing, China). Blood biochemistry, including aspartate aminotransferase (AST), alanine aminotransferase (ALT), alkaline phosphatase (ALP), total bilirubin (TBIL), direct bilirubin (DBIL), total protein (TP), albumin (ALB) were determined using an automated biochemistry analyzer (HITACH 7020, Japan Hitachi Corporation, Tokyo, Japan), and the reagents were purchased in Lepu (Beijing) Diagnostics Co., Ltd. AFB1 residue was analyzed with ELISA kits (Pribolab Biological Engineering Co., Ltd. Qingdao, China).

### 2.4 Statistical analysis

All data were analyzed by one-way analysis of variance (ANOVA) according to significant difference test, using SPSS 26.0 statistical package. Comparisons among groups were performed using Duncan’s multiple comparisons test. Results were expressed as mean value and the standard error of the mean (SEM). Data means significance was declared at P ≤ 0.05 and tendencies were considered at 0.05 < P < 0.1.

## Author Contributions

Conceptualization, H.R.W.; Formal Analysis, Y.Z. and H.N.L.; Investigation, Y.Z. and H.N.L.; Data Curation, Y.Z. and H.N.L.; Writing—Original Draft Preparation, Y.Z. and H.N.L.; Writing—Review and Editing, E.K. and H.R.W.; Supervision, H.R.W. All authors have read and agreed to the published version of the manuscript.

## Funding

This research was funded by the project named the Study on Green Control Technology of Mycotoxin in Silage Maize (2022-Scientific Research Tackling -1).

## Institutional Review Board Statement

The animal study protocol was approved by the Inner Mongolia Agricultural University Scientific research and academic ethics special committee for studies involving animals.

## Data Availability Statement

The data are available from the corresponding author on reasonable request.

## Acknowledgments

The authors thank the National Dairy Technology Innovation Center for providing support for this project.

## Conflicts of Interest

The authors declare no conflict of interest.

## Notes

### Competing Interest Statement

The authors have declared no competing interest.

